# High-throughput single-cell DNA methylation and chromatin accessibility co-profiling with SpliCOOL-seq

**DOI:** 10.1101/2025.02.25.640234

**Authors:** Qingmei Shen, Enze Deng, Jingna Zhang, Qifeng Yang, Dan Su, Xiaoying Fan

## Abstract

DNA methylation and chromatin accessibility are fundamental epigenetic mechanisms that orchestrate gene expression programs, define cellular states, and drive developmental trajectories. scCOOL-seq has enabled simultaneously measuring the two modalities in the same single cells, but in quite a low throughput manner. We present single-cell split-pool ligation-based multi-omics sequencing technology (SpliCOOL-seq), which improves the throughput to thousands of cells by combining split-pool ligation based single-cell indexing after in situ tagmentation with universal Tn5 transposase and scCOOL-seq. SpliCOOL-seq achieved higher sensitivity than previous high throughput single-cell DNA methylation sequencing methods and can clearly distinguish different lung cancer cells based on both genetic and multiple epigenetic modalities. We show that the two DNMT inhibitors, 5-Azacitidine and Decitabine, both cause large scale demethylation but in distinct patterns. Applied to the primary lung tumor, SpliCOOL-seq clearly captured subclones within the tumor lesion and revealed candidate genes related to tumorigenesis. Furthermore, we presented the first report on the heterogeneity of scDNAm age acceleration among tumor subclones as predicted from a single-cell perspective. In conclusion, SpliCOOL-seq achieves parallel profiling of whole genome DNA methylation and chromatin accessibility in the same individual cells in a high-throughput manner and is hopefully used to illustrate regulatory interactions under different cell states.

## Introduction

Both chromatin accessibility and DNA methylation are critical regulations controlling cell-type-specific gene expression patterns. DNA methylation, predominantly occurring at CpG dinucleotides, mediates transcriptional silencing by hindering transcription factor binding or recruiting of methyl-binding proteins[1–3]. Chromatin accessibility, reflecting nucleosome positioning and the activity of regulatory elements, determines the spatial availability of DNA for transcriptional machinery and enhancer-promoter interactions[4]. The interplay between these two epigenetic modalities is critical for maintaining cellular homeostasis. For example, global DNA methylation erasure in primordial germ cells coincides with chromatin remodeling, resetting pluripotency[5]. Additionally, hypomethylation of oncogenic promoters and hypermethylation of tumor suppressor genes are often coupled with chromatin compaction at regulatory regions[6,7]. Thus, simultaneous detection and integrative analysis of whole genome DNA methylation landscape alongside chromatin accessibility can provide a comprehensive understanding of how these different epigenetic modalities coordinate to establish and maintain cellular identity, as well as the critical changes that occur in disease[8–14].

The scCOOL-seq[8] enables concurrent measurement of multiple epigenetic modalities within a single cell, leading to subsequent development of related multi-omics methodologies including scNOME-seq[9], scNMT-seq[10], iscCOOL-seq[11], scChaRM-seq[12], snmCAT-seq[13], and scNOMeRe-seq[14]. However, the utilization of these multi-omics techniques has been restricted to single-cell lysates, which severely limited the cell throughput and incurs substantial costs and time consumption. Achieving high-throughput single-cell sequencing that can simultaneously detect genome-wide DNA methylation and chromatin accessibility remains a challenge.

In this study, we introduce SpliCOOL-seq, an improved method that combines split-pool ligation-based single-cell indexing following in situ GpC methylation and tagmentation using universal Tn5 transposase. This innovative approach enables high-throughput analysis of nucleosome-depleted regions (NDRs), also known as chromatin accessible regions, alongside genome-wide DNA methylation. SpliCOOL-seq demonstrates superior read quality and sensitivity compared to previous high throughput single-cell DNA methylation sequencing approaches, allowing for precise discrimination among heterogeneous lung carcinoma populations based on genomic and multiple epigenetic features. Our findings reveal that different DNMT inhibitors induce extensive demethylation in distinct manners, suggesting that the choice of drugs for clinical cancer treatment should be carefully evaluated in advance. Furthermore, we identified subclones within a lung tumor lesion that exhibited distinct genomic and epigenomic characteristics. Overall, SpliCOOL-seq represents a significant advancement in the repertoire of multimodal single-cell profiling technologies, proving to be a valuable tool for atlas studies.

## Results

### SpliCOOL-seq enables co-profiling of DNA methylation and NDRs in a high throughput manner

We presented a strategy that enables the concurrent assessment of genome-wide DNA methylation (WCG) and chromatin accessibility (NDRs, via GCH methylation) in individual cells (Fig. 1A). Initially, *in situ* methylation of GpC sites was performed while maintaining the chromatin structure. The fixation of cell nuclei is critical and should be occur after GpC methylation treatment using an appropriate concentration of formaldehyde (Additional file 1: Supplementary Fig. 1A-C). Unlike previous single-molecule combinatorial indexing (sci)-based methods[15–17], we utilized universal Tn5 transposase complexes. This approach was adopted to avoid variations in Tn5 transposase activity and fragmentation efficiency that may arise from incorporating Tn5 transposase with different barcodes within each reaction chamber. As a result, we achieved more uniform data quantity across cells compared to sciMETv2[17] (Additional file 1: Supplementary Fig. 1D). Published single-cell whole genome bisulfite sequencing methods utilizing post-bisulfite adapter-tagging (PBAT) often exhibit suboptimal mapping efficiency[18], likely due to substantial generation of chimera fragments during the random priming step. We observed larger library sizes and lower mapping ratios when conducting four rounds of random priming and extension (Additional file 1: Supplementary Fig. 1E, F); however, this did not adversely affect genome coverage (Additional file 1: Supplementary Fig. 1G). The implementation of fluorescence activated cell sorting (FACS) in the final splitting step has the potential to reduce sequencing background, but no significant difference was observed in the read ratio within cells (∼90%) (Additional file 1: Supplementary Fig. 1H, I).

**Figure 1.**
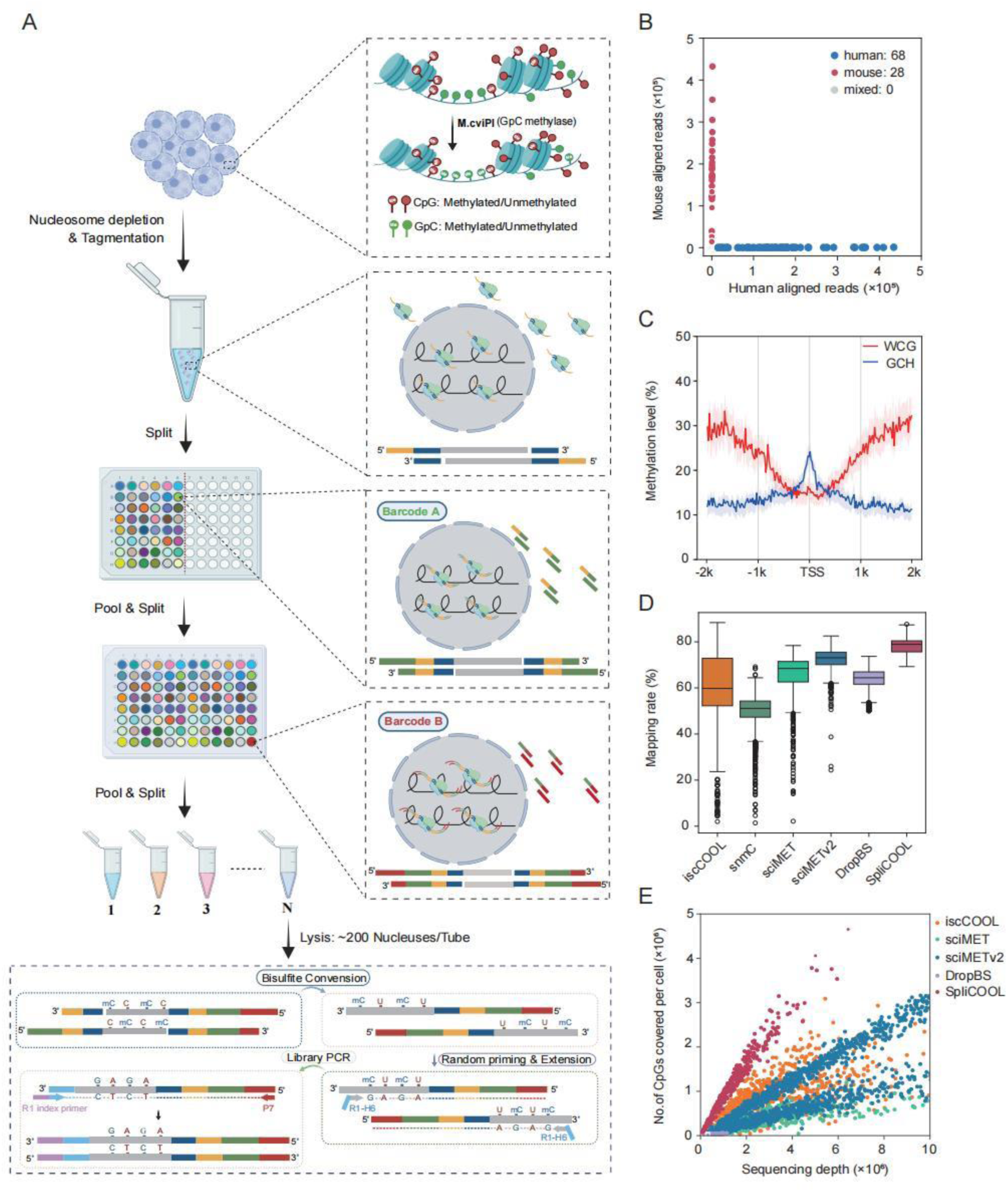
SpliCOOL-seq overview and performance. **A)** Schematic of SpliCOOL-seq. Single nuclei were treated with M.CviPI in situ, followed by fixation and disruption of the nucleosomes. Then nuclei were barcoded through split-pool combinatorial indexing after universal Tn5 tagmentation. The nuclei were subsequently lysed, followed by bisulfite conversion, random priming and extension. The final library was obtained through PCR amplification. **B)** Scatter plot illustrating the number of reads mapped to the human and mouse genomes in each cell from the species-mixing experiment. **C)** WCG and GCH methylation levels of GM12878 cells within ±2 kb of the transcription start site (TSS) as determined by SpliCOOL-seq, with shaded regions indicating the 25th and 75th percentiles of methylation levels across cells. Cell number n=84. **D)** Comparison of mapping efficiencies across individual cells by each method. **E)** Scatter plot depicting the number of CG dinucleotides covered by the total aligned reads per cell by each method.

Species-mixing experiments revealed a negative correlation between the number of cells per tube in the final splitting step and the anticipated collision rate. When the cell count per tube was below 100, doublets were virtually absent, while the expected collision rate was approximately 13.5% for barcodes with ∼300 cells per tube (Fig. 1B and Additional file 1: Supplementary Fig. 1B). Consistent with previous studies[9,10], SpliCOOL-seq detected gene transcription start sites (TSS) with DNA hypomethylation and enhanced accessibility (Fig. 1C). In comparison to earlier single-cell DNA methylation sequencing techniques[11,16,17,19,20], SpliCOOL-seq achieved a superior read mapping rate (from 69.3% to 87.6%) and detected a greater number of CpG sites at equivalent sequencing depths within individual cells (Fig. 1D, E), indicating high read quality and sensitivity. Notably, at the resolution of single WCG sites, approximately 100 cells can achieve 80% (≥1x depth) and 70% (≥5x depth) genomic coverage, respectively (Additional file 1: Supplementary Fig. 1J). Collectively, our findings demonstrate that SpliCOOL-seq proficiently captures both DNA methylation status and chromatin accessibility at single-cell level in a high throughput manner.

### SpliCOOL-seq clearly distinguishes different lung cancer cells at multiple omics dimensions

To evaluate the capability of SpliCOOL-seq in resolving cell-type-specific DNA methylation and chromatin accessibility, we applied the technique to three lung cancer cell lines including A549, H460 and SK-MES. A total of 1,310 cells were obtained, yielding an average of 463,522 unique reads, 193,554 WCG sites and 1,689,066 GCH sites per cell (Fig. 2A). We assessed the DNA methylation levels (WCG%) within gene regions and chromatin accessibility (GCH%) around the transcription start sites (TSS) for each cancer cell type (Fig. 2B). All cells exhibited dropped WCG methylation levels at the TSS, while the gene body showed the highest WCG methylation compared to flanking regions. Although A549 cells displayed lower genome-wide WCG methylation levels, they exhibited significantly higher methylation around gene-franking regions (Fig. 2A, B), indicating the importance of evaluating DNA methylation levels in distinct genome contexts. Furthermore, lower levels of DNA methylation did not correspond to higher chromatin accessibility in different cell types (Fig. 2B), underscoring the complexity of epigenetic co-regulation.

**Fig 2.**
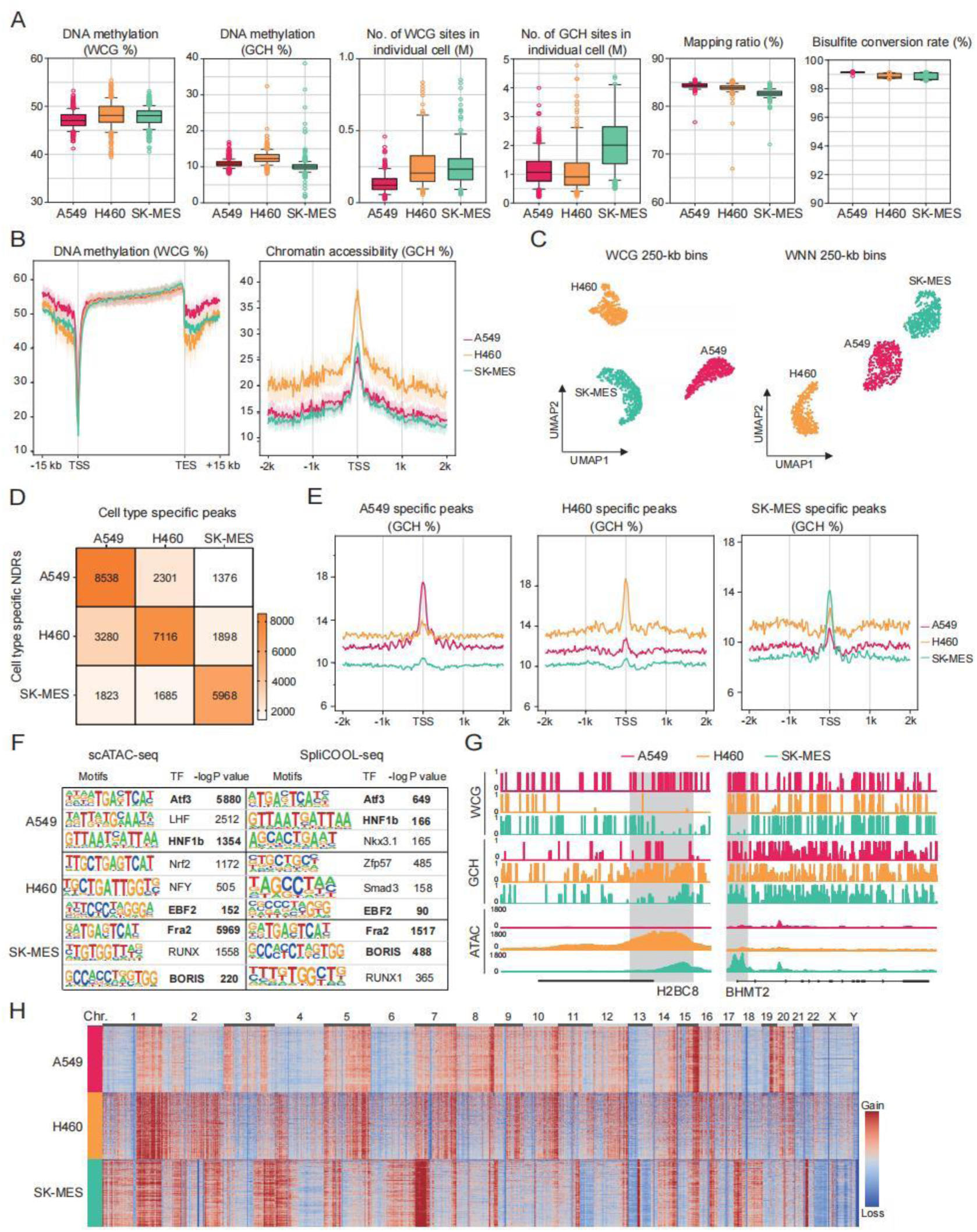
SpliCOOL-seq distinguishes different cell types based on multiple modalities. **A)** Global DNA methylation levels (as measured by the modification levels of WCG%), chromatin accessibility (as measured by the modification levels of GCH%), the numbers of WCG sites and GCH sites covered, mapping ratio, and bisulfite conversion rate estimated in individual cells of A549 cells, H460 cells and SK-MES cells profiled by SpliCOOL-seq. **B)** DNA methylation levels within and surrounding gene bodies (left panel) and chromatin accessibility around the TSS (right panel) in single cells belonging to the three cell types. Solid lines represent the mean levels of single cells, while shaded areas indicate the 25th and 75th percentiles among the cells. **C)** UMAP embedding showing cells based on their DNA methylation levels (WCG in 250-kb non-overlapping bins) and an integrated analysis of DNA methylation and chromatin accessibility (WNN in 250-kb non-overlapping bins). **D)** Heatmap showing the intersection of cell-type-specific peaks identified through scATAC-seq and NDRs detected via SpliCOOL-seq. The corresponding numbers of overlapped regions are displayed within the boxes. **E)** Chromatin accessibility around the TSS which were cell type specific based on scATAC-seq analysis. **F)** Top enriched motifs for cell-type-specific accessible regions and NDRs are identified from different methods. P values are calculated using one-sided Fisher’s exact test. **G)** Browser track showing the DNA methylation (WCG) and chromatin accessibility (GCH or ATAC) profiles at the *H2BC8* locus and *BHMT2* locus across various cell types. Promoter regions are highlighted by light gray shading. **H)** Heat map showing the CNVs in individual A549, H460 and SK-MES cells at 10 Mb resolution, as inferred from SpliCOOL-seq data.

By partitioning the genome into 250-kb bins, both WCG and GCH methylation data distinctly segregated the three cell types into separate clusters through unsupervised clustering (Fig. 2C and Additional file 1: Supplementary Fig.2A). Integration of the two modalities using weighted-nearest neighbor (WNN)[21] yielded comparable clustering outcomes (Fig. 2C). At the current sample size, DNA methylation was more effective at distinguishing different cancer cell types, with TSS methylation data alone sufficient to separate the cells in the uniform manifold approximation and projection (UMAP) analysis (Fig. 2C and Additional file 1: Supplementary Fig.2B). To conform that the NDRs we profiled represent true accessible chromatin regions, we also performed scATAC-seq analysis on the same cells (Supplementary Fig. 2C, D). The cell type-specific NDRs and differentially accessible regions (DARs) showed substantial overlap (Fig. 2D, Additional file 2: Table S1). The DARs for each cell type also showed higher GCH methylation levels in the corresponding cell type (Fig. 2E). Notably, the enriched motifs identified in both spliCOOL-seq and scATAC-seq were identical in each cell type (Fig. 2F), suggesting that SpliCOOL-seq accurately captures open chromatin regions.

We then calculated the differential expressed genes (DEGs) across the cell types and found cell-type specific DEGs (Additional file 2: Table S1). For instance, we observed elevated GCH methylation levels of *H2BC8* and *H2AC8* in H460, which corresponded to peak-rich regions in the scATAC-seq analysis, while the WCG methylation levels at TSS of these genes remained significantly lower (Fig. 2G and Additional file 1: Supplementary Fig. 2E-G). Similarly, betaine-homocysteine S-methyltransferase2 (*BHMT2*), a marker for lung squamous cell carcinoma (LUSC) cell line SK-MES[22], exhibited higher GCH methylation levels and lower WCG methylation levels in SK-MES cells, indicating a negative regulatory relationship between the two modality at gene regions. We further categorized the genes into four groups based on their expression levels within each cell type (using ENCODE data; see methods). The promoter methylation levels gradually decreased while promoter accessibility increased in tandem with elevated gene expression (Additional file 1: Supplementary Fig. 2H), consistent with observations from the previous study[23]. Thus, SpliCOOL-seq enables integrated analysis of DNA methylation and chromatin accessibility within the same single cells.

Since scCOOL-seq also facilitates the analysis of copy number variations (CNVs), we assessed the ability of SpliCOOL-seq to capture CNVs in different cancer cells. Although we observed reduced genome coverage compared to traditional scCOOL-seq, SpliCOOL-seq accurately captured the CNVs in each cancer cell types, allowing differentiation of various tumor clones (Fig. 2H and Additional file 1: Supplementary Fig. 2I). It is worth mentioning that the scATAC-seq data could not specify CNVs in different cancer cells using the same bioinformatics pipeline (data not shown, see methods), highlighting the superiority of SpliCOOL-seq in capturing genomic features.

### DNMT inhibitors exhibit distinct patterns in the demethylation of tumor cells

Currently, DNA demethylation agents, also known as DNMT inhibitors, such as azacitidine (5-Aza) and decitabine (DEC), exert their therapeutic effects by interfering with the DNA methylation machinery[24]. These drugs have shown significant efficacy in treating myelodysplastic syndromes[25], acute myeloid leukemia[26], and other malignancies that are refractory to conventional chemotherapy[27]. To investigate the effects of these agents, we treated A549 cells with 5-Aza or DEC for 1, 3 and 5 days and evaluated the changes in DNA methylation and chromatin accessibility using SpliCOOL-seq (Fig. 3A). The global methylation level exhibited a gradual decline with extended treatment. By day 5, the averaged global DNA methylation level decreased from 48.03% to 32.94% in the 5-Aza group and to 25.13% in the DEC group (Fig. 3B). However, chromatin accessibility did not show corresponding changes (Additional file 1: Supplementary Fig. 3A). Thus, the reduction in methylation levels induced by these hypomethylating agents did not elicit corresponding alterations in chromatin status *in vitro*[28].

**Fig 3.**
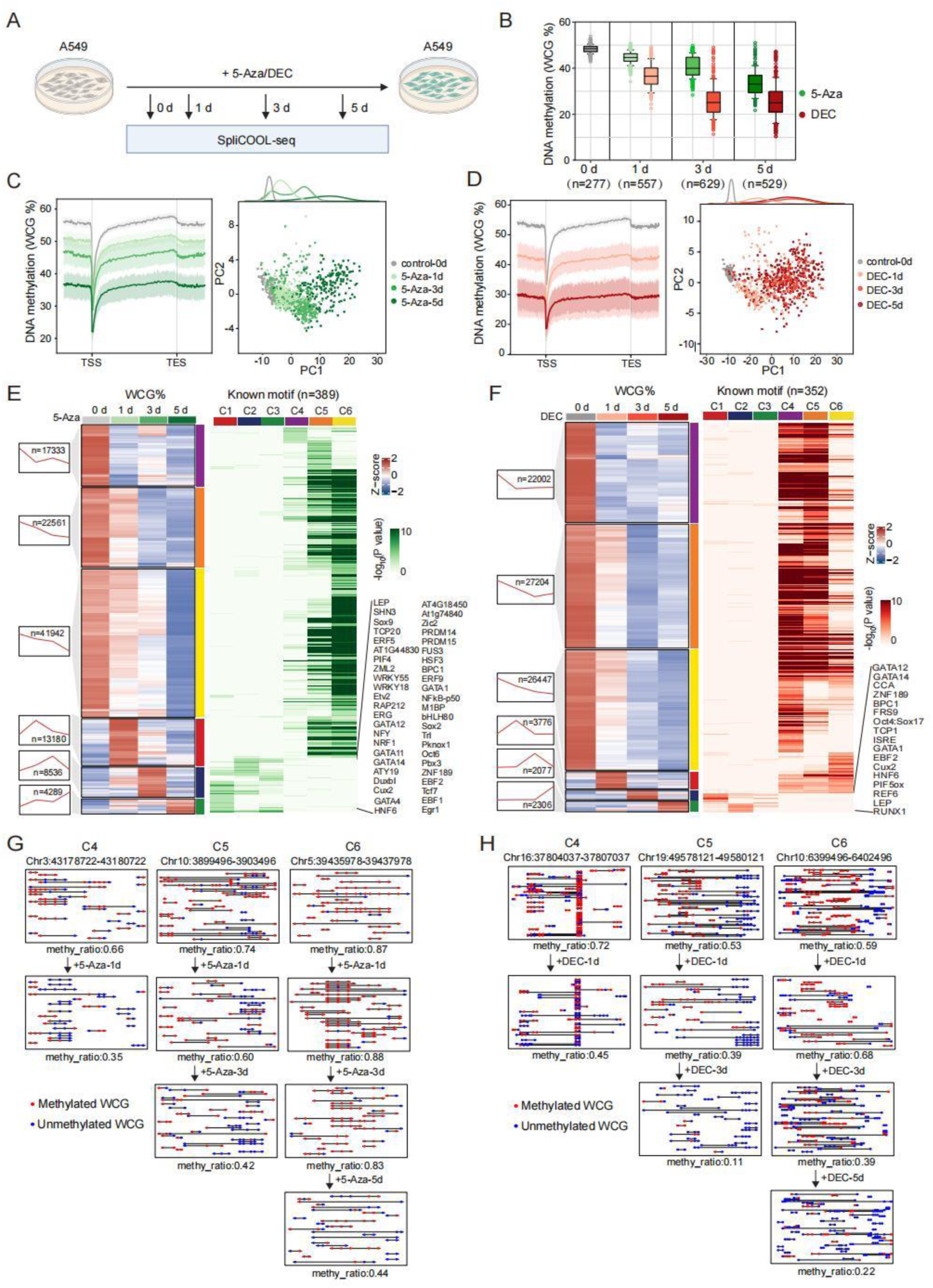
SpliCOOL-seq detected distinct demethylation patterns of tumor cells under treatment with different DNMT inhibitors. **A)** Schematic illustration of the treatment protocol for A549 cells with 5-Aza or DEC. Cells are subjected to 1 μM concentrations of 5-Aza or DEC for durations of 0, 1, 3, and 5 days. **B)** Boxplot showing global WCG methylation levels in cells treated with 5-Aza (green) or DEC (red) at each timepoint. **C, D)** Average WCG methylation levels across the gene body in single cells after being treated with 5-Aza (**C**) and DEC (**D**) for 0, 1, 3, and 5 days. 15 kb upstream the TSS and 15 kb downstream the TES were calculated (Left). The shading areas display the 25th and 75th percentile of DNA methylation levels across cells. The right panels display Principal component analysis (PCA) of all the cells based on WCG in 250-kb non-overlapping bins. **E, F)** heatmaps showing the DMR clusters showing different methylating variations along time. The 6 clusters are represented in distinct colors on the right side of the heatmap (left). The enriched motifs in each cluster are shown in the right panels, with the TF lists of C1-C3 clusters presented. **G, H)** The three clusters of DNA demethylation patterns in A549 cells in response to the 5-Aza treatment (**G**) or DEC (**H**) treatment. Red circles and blue circles represent methylated and unmethylated WCG sites, respectively.

We observed a marked difference in the demethylation rhythms between the two agents. 5-Aza induced a gradual decrease in methylation levels over time, both genome-wide and in gene regions, while DEC treatment did not result in significant changes from day3 to day5 (Fig. 3B-D). Similar demethylation patterns were observed across other genomic elements (Additional file 1: Supplementary Fig. 3B, C). To further investigate the similarities and differences in the demethylation processes induced by the two agents, we analyzed differential methylation sites (DMSs). 5-Aza induced more DMSs than DEC, but the magnitude of demethylation changes was smaller in the early stages (Additional file 1: Supplementary Fig. 3D), indicating that 5-Aza results in greater heterogeneity across cells. Although both groups exhibited tens of thousands of hypomethylated DMSs between adjacent stages, there were also thousands of hypermethylated DMSs (Additional file 1: Supplementary Fig. 3D). Notably, from day 3 to day 5 in the DEC group, the number and magnitude of hypermethylated and hypomethylated DMSs were nearly equal, resulting in globally unchanged methylation levels (Fig. 3D, and Additional file 1: Supplementary Fig. 3D). In both groups, the hypomethylated DMSs were specific at each time point, suggesting that the demethylation process was sequentially happened for all the target sites (Additional file 1: Supplementary Fig. 3E, F).

We also performed clustering of differentially methylated regions (DMRs) in each group, which could be divided into 6 clusters based on methylation changes over time (Fig. 3E, F, and Additional file 3: Table S2). Consistent with the DMS analysis, the majority of DMRs underwent demethylation (C4-C6 clusters), while a subset of DMRs transiently gained methylation (C1-C3 clusters). Specifically, the methylation level of C4 DMRs dropped to its lowest on the first day of treatment, suggesting these regions might be directly targeted by DNMT. The C5 and C6 DMRs were later demethylated, indicating they were indirectly regulated by DNMT (Fig. 3E-H). The three transiently hypermethylated DMRs were enriched in similar motifs, which were also largely shared between the two agent treatment groups, such as GATA1/12/14, LEP, ZNF189 and BPC1 (Fig. 3E, F). Further gene ontology (GO) analysis of these DMR-related genes revealed that both 5-Aza and DEC induced similar gene regulatory programs. For instance, C4-C6 genes were enriched in developmental-related processes, cilium assembly and response to stimulus; while cell proliferation, G protein-coupled receptor activity and nervous system processes related genes exhibited transient gains in methylation (C1-C3) in both groups (Additional file 1: Supplementary Fig. 3G, H). Additionally, the two DMNT inhibitors induced specific regulatory changes. For example, chromatin remodeling and nuclear body organization genes were specifically demethylated in the DEC group, while biosynthetic processes related genes were affected only in the 5-Aza group (Additional file 1: Supplementary Fig. 3G, H). Together, these results indicate that although different DNMT inhibitors induce global demethylation in cancer cells, they exhibit distinct preferences for target regions, which should be considered in clinical practice.

### SpliCOOL-seq revealed tumor subclones within the tumor lesion

Due to the limitations of current single-cell methylation sequencing techniques, the genomic and epigenetic heterogeneity of primary cancer cells has been insufficiently characterized. In this study, we applied SpliCOOL-seq for an integrated analysis of genomic alterations, aberrant DNA methylation and chromatin accessibility in a lung adenocarcinoma (LUAD) tissue sample. The tissue was initially divided into two parts—the primary tumor (PT) and adjacent tissue (AT) (Fig. 4A). We obtained a total of 840 cells from the AT, with an average of 298,447 unique reads per cell, and 777 cells from the PT, with an average of 370,702 unique reads per cell (Fig. 4A and Additional file 1: Supplementary Fig. 4A). Notably, because we extracted the nuclei from the tissues for SpliCOOL-seq (see methods), the majority of the cells from both AT and PT were epithelial cells, which exhibited specific hypomethylation and chromatin accessibility in the marker gene EPCAM (Fig. 4B and Additional file 1: Supplementary Fig. 4B). All single cells could be clustered into two groups based on both DNA methylation levels (using WCG 250-kp bins) and chromatin accessibility (based on NDRs) (Fig. 4C). The two cell groups were quite matched in both modalities, demonstrating region specificity, although a few PT cells clustered with those from the AT (Fig. 4C and Additional file 1: Supplementary Fig. 4C). Further in-depth examination of somatic CNVs[28] revealed that all cells could be categorized into four types: primary cancer cells, which were further subdivided into tumor subclone PC1 and PC2, exhibiting a high abundance of somatic CNVs; adjacent normal (AN) cells, which displayed a normal diplotype; and adjacent cancerous (AC) cells, which harbored a small number of somatic CNVs (Fig. 4D). Interestingly, the AC cells were dispersed across both AT and PT regions and exhibited DNA methylation and chromatin accessibility patterns more similar to AN cells (Additional file 1: Supplementary Fig. 4C), suggesting that these AC cells may represent early cancerous cells.

**Fig 4.**
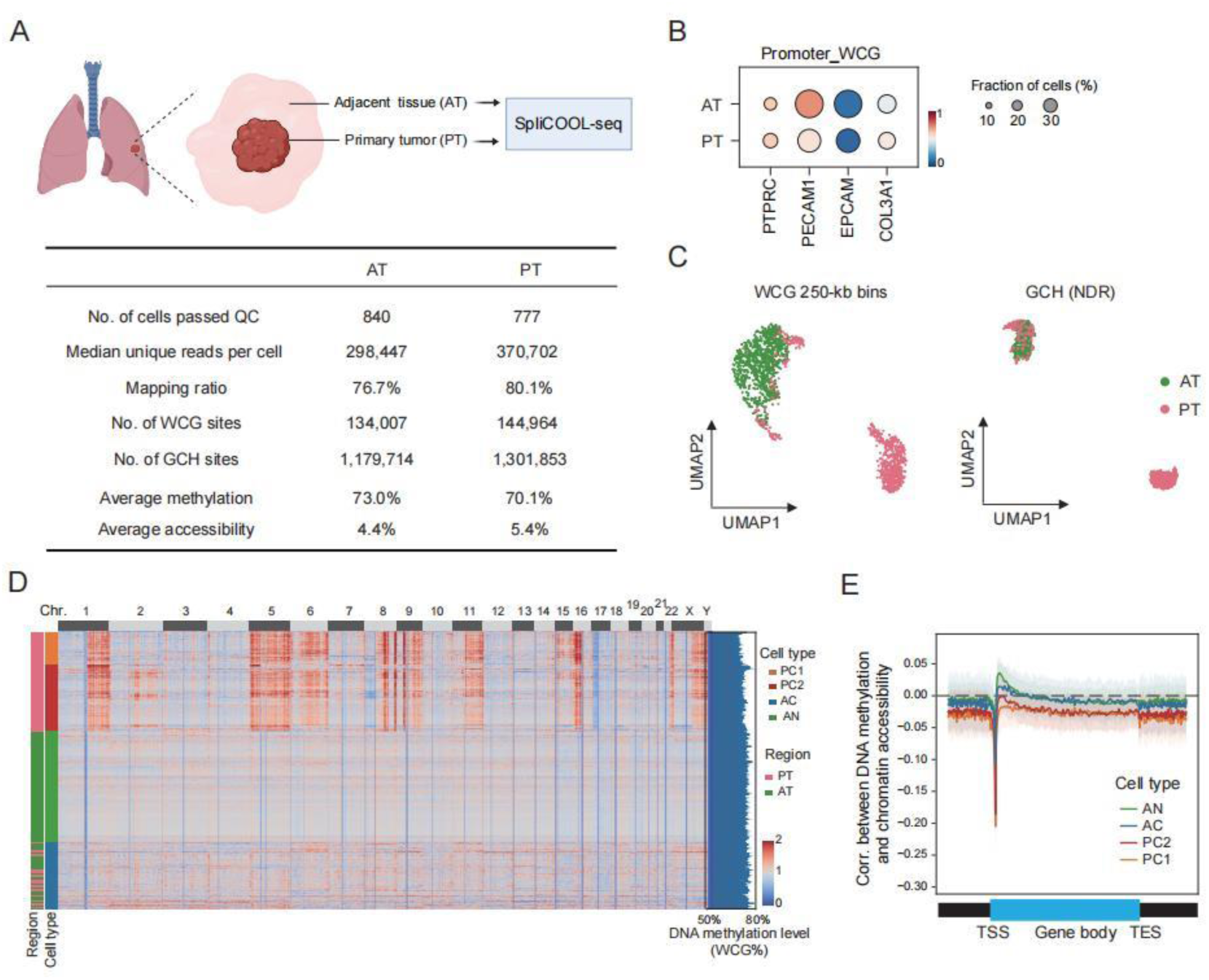
SpliCOOL-seq revealed tumor subclones within the tumor lesion. **A)** Schematic illustration of sampling (upper). The lower table presents the data quality of AT and PT cells in the current experiment. **B)** Dot plots showing the promoter WCG methylation levels (−1,000 bp to +1,000 bp) of representative marker genes in each tissue section. **C)** UMAP embedding showing cells based on their DNA methylation levels (WCG in 250-kb non-overlapping bins) and chromatin accessibility (GCH based on NDRs). **D)** Heatmap showing the high frequency of subchromosome-scale CNV pattern in PC cells. Bar plot on the right shows the global DNA methylation level of each individual cell. **E)** Spearman correlations between DNA methylation (WCG%) and chromatin accessibility (GCH%) across the gene body in each cell type.

We further analyzed the changes in the three clones of cancerous cells compared to AN cells. The global methylation levels of the two PC clones were significantly decreased (averagely 68.2% for PC1 and 70.1% for PC2) compared to that of AN cells (73%), while the methylation levels of AC cells remained relatively unchanged (72.6%, Additional file 1: Supplementary Fig. 4D, E). Moreover, the interaction between DNA methylation and chromatin accessibility was maintained in AC cells, whereas the PC cells showed a stronger negative correlation between these two modalities (Fig. 4E). Together, we captured a group of early cancerous cells that display genomic abnormalities without significant changes in their epigenome.

### SpliCOOL-seq enables the identification of candidate tumor diagnostic targets

With the identification of tumor subclones, we can capture molecular targets associated with tumor progression that may have been masked in whole population analyses. Our analysis revealed that PC1 cells had the highest number of methylation aberrant genes (MAGs) compared to AN cells, with over 60% of the MAGs in PC2 cells overlapping with those in PC1 (Fig. 5A, Additional file 1: Supplementary Fig. 5A and Additional file 4: Table S3). In contrast, only 422 MAGs were observed in AC cells, and less than 20% shared with the PC cells, indicating that AC cells may be independently derived. Furthermore, GO analysis demonstrated that the hypomethylated genes in AC cells were significantly enriched in multiple catabolic processes, which markedly differed from the enrichment profiles in PC1 and PC2 cells, both of which were related to sensory perception of smell (Additional file 1: Supplementary Fig. 5B). Interestingly, the hypermethylated genes in AC cells shared pathways with those in the PC cells, suggesting that early tumorigenesis may be accompanied by abnormal translation processes (Additional file 1: Supplementary Fig. 5C).

**Fig 5.**
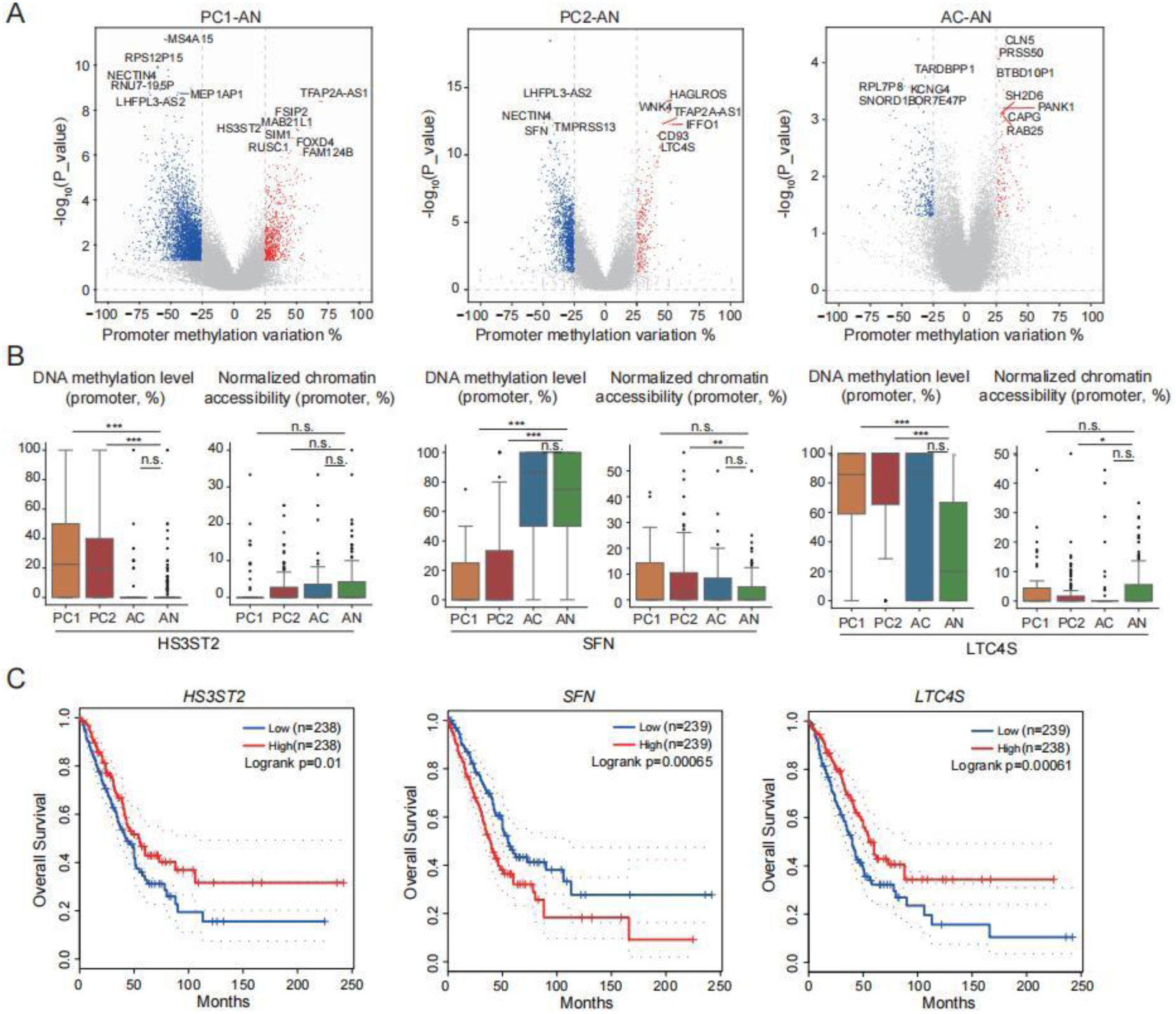
SpliCOOL-seq enables identification of candidate tumor diagnostic targets. **A)** Gene promoter methylation variations between each cancerous clone and AN cells. Top representative genes within each clone are labeled. **B)** DNA methylation levels and chromatin accessibility in the promoter regions of the candidate biomarkers across each cell type. The statistical test was carried out using the Wilcoxon rank-sum test. n.s., no significance; *P < 0.05; **P < 0.01; ***P < 0.001. **C)** Overall survival of LUAD patients grouped by the expression of the representative genes.

The top differentially methylated genes identified across the three cancerous clones were highlighted in Fig. 5A and may serve as potential DNA methylation biomarkers for LUAD. For example, *HS3ST2* and *LTC4S*, both previously reported in non-small cell lung cancers (NSCLCs)[29,30], were significantly hypermethylated in the two PC clones, and higher expression of these genes was associated with improved overall survival in LUAD (Fig. 5B, C). Additionally, we identified the hypomethylated gene *SFN* in PC cells, whose overexpression has been significantly linked to gallbladder cancer[31] and cervical cancer[32]. *SFN* also showed higher chromatin accessibility in PC2 cells (Fig. 5B), suggesting it may be a novel therapeutic target for LUAD. The AC cell-specific hypomethylated gene *OR7E47P* exhibited contrasting overall survival in LUAD and in lung squamous cell carcinoma (LUSC), but in reversed patterns (Additional file 1: Supplementary Fig. 5D). Previous studies have identified *OR7E47P* as having ubiquitous expression in lung tissue and a positive correlation with apoptosis[33]. It has also been characterized as a positive regulator within the tumor microenvironment of LUAD[34]. Consequently, *OR7E47P* may serve as a potential biomarker for the early detection and screening of LUAD.

We further investigated the hypermethylated genes *HS3ST2* and *LTC4S* in various lung cancer cell lines and confirmed their hypermethylated status specifically in the LUAD cell line A549. Following treatment with two hypomethylating agents, the two genes exhibited distinct demethylation patterns. Notably, both genes underwent gradual demethylation with 5-Aza, while DEC induced only transient hypomethylation of *HS3ST2* at early stages (Additional file 1: Supplementary Fig. 5E). Moreover, DEC elicited a more pronounced demethylation effect on *LTC4S*. These findings reinforce the potential of SpliCOOL-seq in identifying novel cancer biomarkers and highlight the functional distinctions between different demethylation drugs.

### Cancer cells exhibit accelerated epigenetic aging

The development of epigenetic clocks to predict physiological age based on DNA methylation has become a pivotal resource for comparative epigenomic studies, shedding light on the mechanisms underlying epigenetic aging[35–37]. We sought to determine whether SpliCOOL-seq could capture changes in aging status during tumorigenesis. To this end, we adopted scAge[38] to calculate the epigenetic age of individual cells. First, we utilized the 450k methylation chip data of LUAD downloaded from The Cancer Genome Atlas (TCGA) for model training, which also returned the rank of CG sites correlated with age (Additional file 5: Table S4). For the SpliCOOL-seq covered sites in each single cell, we found only about 10% were overlapped with those in the chip across different clones (Additional file 1: Supplementary Fig. 6A, B), indicating that SpliCOOL-seq captures CG sites which are largely missed by methylation chip assays. Then, the top 10% aging associated CG sites in each clone were used to predict the single-cell DNA methylation (scDNAm) age. We compared these sites were with the CpG markers reported in existing epigenetic clocks[39–49], finding they were significant overlapped with the markers in three canonical DNA methylation-based epigenetic clocks – Hannum[42], Yang[46] and Horvath2[48] (Fig. 6A). This convergence underscores the evolutionary conservation of epigenetic regulatory frameworks and validates the predictive accuracy of our model.

**Fig 6.**
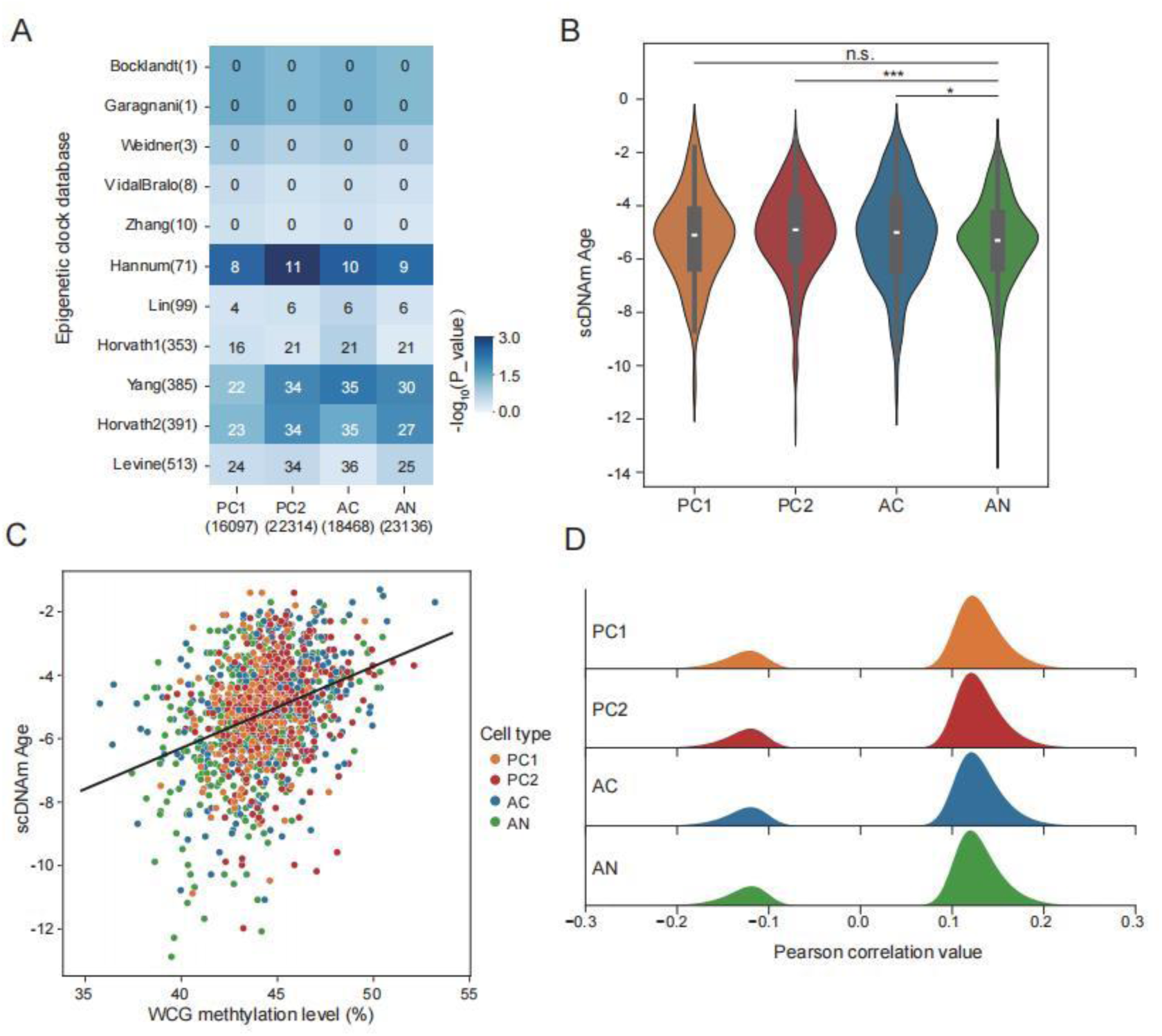
Cancer cells exhibit accelerated aging. **A)** Heatmap showing the intersection of top 10% age-associated CG sites identified in each clone and those identified in different epigenetic clocks. The numbers of CG sites are presented within the respective brackets. **B)** Violin plots depicting the scDNAm Age distribution in each clone. **C)** Dot plot illustrating the correlation between global methylation levels of the predictive CG sites and scDNAm Age for each cell, with point colors representing distinct clones. **D)** Distribution of pearson correlation values in the scAge model for the predictive CG sites in each clone.

Based on the in home trained model, we found that nearly all three cancerous clones showed increased epigenetic age compared to the AN cells (Fig. 6B), consistent with previous findings that cancer tissues exhibit significant age acceleration[40,42,46]. Analyzing the relationship between the methylation levels of the predictive CG sites and the scDNAm age revealed a positive correlation across all cell clones (Fig. 6C), suggesting that the majority of the aging associated sites gained methylation along time, which even intense in LUAD cancer cells. Further dividing of the predictive sites based on their correlation with age showed that 80.3% sites showed age-positive correlation in both tumor and normal cell clones (Fig. 6D). These findings indicate that aging and LUAD tumorigenesis may share similar epigenetic signals, a phenomenon also noted in other types of cancer[50–52].

To specify how tumorigenesis affects epigenetic aging, we extracted sites positively correlated with age that also showed elevated methylation level in tumor cells. We identified a total of 124 sites from all three cancerous clones, which corresponded to 138 genes (Additional file 6: Table S5). GO analysis of these genes showed significant enrichment in calcium ion homeostasis, organ morphogenesis and neuronal system (Additional file 1: Supplementary Fig. 6C), which have been previously reported to be associated with cellular senescence or tumorigenesis[53–57]. Specifically, *VSIG4* identified in PC1 cells has been identified as a biomarker for adipose tissue macrophages in aged mice. Meanwhile, its expression was markedly induced in the tumor-adjacent stroma in both spontaneous and xenograft models of lung cancer[58]. *IL17C* and *CCL11* (eotaxin) identified in PC2 cells exhibited a significant correlation with age, tumor stage, metastasis, lesion count, and tumor burden[59,60]. *NAF1* identified in AC cells has been reported to modulate calcium ion concentration, reactive oxygen species (ROS) levels, and iron metabolism signaling pathways to fulfill its regulatory functions[61]. Together, we identified a series of abnormal DNA methylation regulations that are shared in aging and tumorigenesis.

## Discussion

We developed SliCOOL-seq, a high-throughput single-cell sequencing technology that enables simultaneous profiling of whole-genome DNA methylation and chromatin accessibility (NDRs). Applied to lung cancer models, this method resolved distinct epigenetic and genomic features across cell types, revealed divergent demethylation patterns induced by DNMT inhibitors, identified tumor subclones with early genomic aberrations, and uncovered heterogeneity in epigenetic age acceleration. These findings highlight SpliCOOL-seq as a powerful tool for multimodal epigenomic analysis.

Recently, version 3 of sciMET[62] has been reported, which also supported the co-profiling of DNA methylation and chromatin accessibility by employing another round of indexed Tn5 tagmentation before the intermediate nucleosome disruption step. Similar to sciMETv2, sciMETv3 conducted the initial round of cell labeling using Tn5 transposase labeled with different indexes, which also led to significant variations for fragmentation among individual cells. Secondly, the nucleosome disruption step utilizing SDS treatment is highly detrimental to the nucleus[63], pre-disruption of the DNA by ATAC tagmentation leads to more severe damage to the nuclear structure, leakage of a large number of free DNA fragments, finally results in increased cross-contamination between cells in the sequencing data. Thus we noticed that there were no cell type-specific peaks in the ATAC data, and there was significantly increased doublets rate in the DNA methylation data in the previous study[62]. Another defect of the previous method is that pen regions make up only a small portion of the genome, thus when co-profiling by different tags, very limited data could be allocated to the ATAC module for each cell, which leads to inefficiency in calling peaks. Of course, SpliCOOL-seq dose present certain limitations. Given that the endogenous CH methylation is a well character in the nervous system, GCH methylation may serve as an imprecise indicator of chromatin accessibility when applied in these contexts. Consequently, this technique may have limited applicability in neurological researches. SpliCOOL-seq uncovered adjacent cancerous cells within the LUAD tissue, which harbored somatic CNVs but retained near-normal DNA methylation levels. These AC cells, dispersed across tumor and adjacent regions, may represent early malignant transformations preceding epigenomic dysregulation. This finding underscores the importance of integrating genomic and epigenomic data to capture incipient tumorigenesis. Moreover, the hypomethylated gene *OR7E47P* in AC cells correlated with improved survival in LUAD, suggesting active anti-tumor processes during the early stages of tumorigenesis. It is worth examining whether enhancing these pathways prevents tumor development.

In this study, we presented the first report on the heterogeneity of scDNAm age acceleration among tumor subclones as predicted from a single-cell perspective. While SpliCOOL-seq revealed accelerated epigenetic aging in tumor subclones, our scDNAm age model was trained on bulk 450k methylation data from TCGA, which differs fundamentally from SpliCOOL-seq profiles in coverage and resolution. This discrepancy may introduce systematic biases for the model. In addition, there were only ∼12% of WCG sites overlapped with the CpGs in the chip, which may also have great impact on the accuracy of the forecast results. Future studies could refine these models using single-cell-specific epigenetic clocks trained on matched datasets to mitigate technical variability.

## Methods

### Cell culture

Cell lines (GM12878, NIH/3T3, A549, NCI-H460, SK-MES) were cultured in 5% CO_2_ at 37 °C. GM12878 and NCI-H460 cells were grown in RPMI1640 (Gibco, C11875500CP) supplemented with 10% (v/v) FBP (HyCyte, C520), 1× penicillin-streptomycin (Gibco, 15140122). NIH/3T3 and SK-MES cells were grown in DMEM (Gibco, C11995500BT), with the same supplement as GM12878 cells. A549 cells were grown in Ham’s F-12K (Gibco, 21127022), with the same supplement as GM12878 cells.

### Cell nuclei isolation

For cultured cells, they were washed with ice-cold 1×PBS buffer (CELLCOOK, CM2018), and spun down at 4 °C and 300g for 5 min. The cell pellet was then re-suspended in ice-cold NIB-HEPES buffer (10 mM HEPES (Sigma-Aldrich, H3375), 10 mM NaCl, 3 mM MgCl_2_, 0.1% Igepal (Sigma-Aldrich, I8896), 1×cOmplete protease inhibitor cocktail, EDTA-free (Roche, 11873580001) and 0.1% Tween (Sigma-Aldrich, P9416)) and incubated on ice for 10 min. Repeated spin down step and re-suspended in ice-cold NIB-HEPES buffer for a second incubation were performed. For clinical tissues, cell nuclei suspensions were prepared by douncing of the liquid nitrogen snap-frozen tissues with NIB-tris Buffer (10mM Tris-HCl pH 7.5 (Sangon Biotech, B548139-0500), 10 mM KCl, 3 mM MgCl_2_, 1×cOmplete protease inhibitor cocktail, EDTA-free (Roche, 11873580001), 0.1% Tween (Sigma-Aldrich, P9416) and 0.1% Igepal (Sigma-Aldrich, I8896)). The nuclei were then washed with ice-cold 1%BSA (Sigma-Aldrich, V900933) after passing through a 40 µm cell strainer.

### In vitro GpC methylation and nucleosome depletion

The single nuclei were washed with 1×DPBS buffer (CORNING, 21-031-CV) and subsequently incubated at 37 °C for 30 min in a reaction mixture containing 1×GpC buffer (New England BioLabs, B0227), 0.6 U/µL M.CviPI methyltransferase (New England BioLabs, M0227L) and 0.6 mM/µL SAM (New England BioLabs, B9003). Then they were washed with ice-cold NIB-HEPES buffer and resuspended in 4ml of NIB-HEPES buffer containing 162µL of 37% formaldehyde (Sigma-Aldrich, F8775), incubated at room temperature for 10 minutes. The fixation was terminated with 0.2M glycine (Sigma-Aldrich, 50046) and incubated on ice for 5 min. After centrifuging at 4 °C and 500g for 5 minutes, the nuclei were washed with 1× NEB buffer 2.1 (New England BioLabs, B7202). Subsequently, the nuclei were resuspended in 1mL of 1× NEB buffer 2.1, followed by the addition of 15ul 20% SDS and incubated at 60 °C for 10 minutes. The single nuclei were washed once with NIB-tris Buffer and then resuspended in 100µL of NIB-tris Buffer.

### Tagmentation and barcoding

The Tn5 transposase complex was prepared by coupling Tn5 enzyme (Vazyme, S111-02) with a universal I7 adaptor (5Phos/GTCTCGTGGGCTCGGAGATGTGTATAAGAGACAG)[64]. The nuclei were distributed into PCR tubes (20,000 nuclei per tube) that contained 0.25µM Tn5 transposase complex, 1×Tn5 tagmentation buffer and were incubated at 55 °C for 30 min with rotation. Then all tagmentated nuclei were pooled together and washed with 1 ml of 1×NEB buffer 3.1 (New England BioLabs, B7203). After centrifuging at 4 °C and 1000g for 5 minutes, the nuclei were resuspended in 478µL of 1×NEB buffer 3.1, followed by the addition of 98.4µL of 10×T4 DNA Ligase Buffer and 24µL of T4 DNA Ligase (ABclonal, RK21500). Next, 12µL ligation mix was distributed to Barcode Plate 01 which contained 8µL of 5µM pre-annealed barcode A primer (Additional file 7: Table S6) and incubated at room temperature for 30 min with rotation. After the first round of ligation, 8µL of 10µM Blocking01-Solution (50 µl of 100 µM Blocker01 oligo, 50 µl of 10× T4 DNA Ligase Buffer and 400 µl of nuclease-free water) was then added to each well, and the blocking reaction was continued at room temperature for 30 min with rotation. The nuclei were pooled together and washed with 1 mL of 1×NEB buffer 3.1. Following centrifuging at 4 °C and 1000g for 5 minutes, the nuclei were resuspended in 956µL of 1× NEB buffer 3.1, supplemented with the addition of 196.8µL of 10×T4 DNA Ligase Buffer and 48µL of T4 DNA Ligase. Subsequently, 12µL ligation mix was dispensed into Barcode Plate02 along with an additional volume of 8µL 5µM pre-annealed barcode B primer. The ligation was performed by incubating the mixture at room temperature for 30 min with rotation. After the second round of ligation, 8µL of 10µM Blocking02-Solution (100 µl of 100 µM Blocker02 oligo, 100 µl of 10× T4 DNA Ligase Buffer and 800 µl of nuclease-free water) was added to each well, and the reaction was continued at room temperature for 30 min with rotation.

### Bisulfite conversion and library preparation

The barcode-ligated nuclei were divided into 200 nuclei per tube and treated with proteinase K (Roche, P8107S), followed by conversion using a Hieff^®^ Superfast DNA Methylation Bisulfite Kit (YESEN, 12225ES50) in accordance with the provided instructions. Each tube was eluted in 20 μL elution buffer and transferred to PCR tubes containing the following reaction mixture for random priming and extension: 2.5 μL of 10×blue buffer (TIANGEN, NG202), 1 μL of 10 mM dNTP mixture (TaKaRa, 4019), and 1 μL of 10 μM random primer (TCGTCGGCAGCGTCAGATGTGTATAAGAGACAGHHHHHH). The reactions were heat-shocked at 95 °C for 45 s and then rapidly cooled on ice for 2 min. Subsequently, each reaction was supplemented with 25 U Klenow (3′5′ exo-) polymerase (TIANGEN, NG202), and incubated at a slow ramp +1 °C /15 s up to 37 °C for 60 min. After random priming and extension, the products were purified with 1× AMPure XP Beads (Beckman, A63882) and eluted with 21 μL of elution buffer. Then they were transferred to a PCR tube containing 25μL of 2× KAPA Hifi HotStart ReadyMix (KAPA BioSystems, KK2602), along with 2 μL of 10 μM QP2 (AACGAGAAGACGGCTACGAGAT) primer and 2 μL of 10 μM N5XX index primer(AATGATACGGCGACCACCGAGATCTACACNNNNNNNNTCGTCGGCAGCGTCAGATGT), following the PCR program: 98 °C for 3 min, 98 °C for 20s, 65 °C for 30s, 72 °C for 30s, step2 to 4 with 12cycles, 72 °C for 5 min and 4 °C hold. The libraries were purified with 0.8× AMPure XP Beads and eluted with 20 μL of elution buffer.

### Library quantification and sequencing

The libraries were quantified using the Qsep100 system to evaluate the size distribution of DNA fragments, while the concentration was determined using Qubit 4.0 fluorometer. Subsequently, these libraries were sequenced on SURFSeq 5000 sequencer in a 150 PE run mode, with read1 125 cycles, read2 125 cycles, i7 index 61 cycles and i5 index 8 cycles.

### scATAC-seq library preparation and sequencing

The three lung cancer cell lines were combined in equal proportions to create a single-nucleus suspension following the protocol outlined in the Chromium Next GEM Single Cell ATAC Reagent Kits v2 (10×Genomics, PN-1000406) User Guide for cell capture and library preparation. The library was subjected to quality control using the Qsep100 system and quantified with the Qubit 4.0 fluorometer. Subsequently, sequencing was performed on the SURFSeq 5000 platform using a 150 PE run mode, with read1 150 cycles, read2 150 cycles, i7 index 8 cycles, and i5 index 16 cycles.

### Pre-processing of SpliCOOL-seq data

The two linker sequences were precisely identified at the fixed position within read 1. Reads lacking these linker sequences were classified as unknown structure. The original barcodes were extracted from the linker sequence and compared against those in the whitelist by calculating the minimum Levenshtein distance. Barcode with a minimum distance exceeding 2 was labeled as unknown barcode and discarded. Trimming and low-quality read filtering were performed using Trim Galore (https://github.com/FelixKrueger/TrimGalore, version 0.6.10) with parameters –length 30 and --quality 20. Reads were then mapped to the mouse GRCm39 or human GRCh38 reference genome using Bismark[65] with Bowtie2[66], applying paired-end and non-directional mapping. After paired-end alignment, unmapped reads were realigned in single-end mode to the same reference genome in single-end mode. PCR duplicates were identified and removed based on their genomic coordinates using the Picard MarkDuplicates tool (https://broadinstitute.github.io/picard/, Version 3.1.1), with the command parameter: “BARCODE_TAG=CB”. Only non-duplicated reads were retained for downstream analysis.

### Quantification of WCG and GCH methylation levels

The methylation levels of cytosines classified as WCGs (W = A or T) and GCHs (H = A, T, or C) with a minimum coverage of one read (≥ 1× depth) were quantified using methylpy[67] with the *call-methylation-state* command. For each WCG or GCH site, the methylation level was calculated as the ratio of reads supporting cytosine (C) to the total number of reads covering the site. The average DNA methylation level and chromatin accessibility for all cells were derived from the mean methylation levels of WCG and GCH sites, respectively. Bisulfite conversion efficiency was assessed by examining the WCH methylation level in mitochondrial DNA. Bigwig files were generated using bed Graph To BigWig (v 2.10)[68].

### Defining NDRs

We identified for single-cell NDRs (scNDRs) in individual cells by adapting methodologies from a prior study[28], employing a custom Python script (from https://github.com/sherryxue-PKU/scNanoCOOL-seq). The detection process can be distilled into three key steps. First, the bam files of a certain group of single cells (either a cell type or a treatment condition) were merged as a pseudo-bulk sample. Then the methylation information at the GCH sites was extracted using the call-methylation-state command from the methylpy software[67]. A sliding-window strategy (with a window length at 100 bp and a step length at 20 bp) was then applied to partition the methylation data of GCH sites into numerous tiles. For each tile, the total number of methylated sites (Cs) and the total number of covered GCH sites (Cs + Ts) were compared to the corresponding chromosomal background using the scipy.stats.chi2 function from the SciPy Python library[69]. Tiles meeting the criteria of a p-value less than 1e-10, coverage of at least one read, and containing more than five GCH sites were designated as NDRs for downstream analysis. Finally, BEDTools[70] was utilized to merge overlapping NDRs, yielding cell type-specific NDRs.

### Processing of RNA-seq data from ENCODE

Raw counts data were obtained from the ENCODE database (ENCFF876SRX, ENCFF147QDQ, ENCFF799PDP, ENCFF328YIW, ENCFF024DUU, ENCFF697YYW, ENCFF477TRE) and the Gene Expression Omnibus (GEO) repository (GSE95536). DEG analysis was conducted using the DESeq function from the DESeq2[71] package, with significance defined by an adjusted p-value threshold of less than 0.05.

### Cell clustering with WCG, GCH and NDR

The WCG and GCH methylation calls were utilized to construct a matrix of cells ×250 kp genomic window. Cells with fewer than 10,000 bins covered and bins with less than 50% of cells coverage were excluded. Missing values in the matrix were imputed using the mean methylation level of each cell. Principal component analysis (PCA) was performed on the matrix to reduce noise, retaining the top 50 principal components for further analysis. A neighborhood graph was constructed to estimate the relationships between data points, and the Leiden algorithm was employed to detect communities and identify cell clusters[72]. Additionally, the same principal components were used for non-linear dimensionality reduction, generating a Uniform Manifold Approximation and Projection (UMAP) visualization.

### scATAC-seq data analysis

The 10x Genomics Cell Ranger-ATAC (v2.1.0)[73] pipeline was executed on the raw sequencing data to generate a count matrix for peaks and transcription factors. The obtained fragments file was loaded and the snapatac2[74] workflow was used for downstream analyses. Cell quality was evaluated based on a minimum TSS enrichment score of 10 and a fragment count range of 5,000 to 100,000 to filter cells. Feature selection was conducted using the pp.select_features function with the parameter n_features=250000. Potential doublet cells were identified and removed by calculating the doublet score for each cell using Scrublet[75]. Following quality control, a total of 5,662 cells were retained for further analysis. Spectral embedding was applied for dimensionality reduction, and UMAP was utilized to project the cells into a two-dimensional space for visualization. Graph-based clustering was performed by constructing a k-nearest neighbor graph using snap.pp.knn, followed by the application of the Leiden community detection algorithm to identify densely connected clusters. Gene activity matrices were computed for each cell using the pp.make_gene_matrix function, and cell type-specific peaks were called using MACS3[76] via the snap.tl.macs3 function.

### Single-cell CNV analysis

The CNVs inferred from the SplicCOOL-seq data were primarily derived from read counts across fixed-length windows using Control-FREEC[77]. The parameters for window size and ploidy were set to 1,000,000 and 2, respectively, in the configuration file. Cells were aggregated into pseudo-bulk profiles by cell type to compute cell type-specific CNVs. The coefficient of variation (CV) for individual cells was determined based on the inferred copy number ratios. Public whole-genome sequencing (WGS) data were acquired from the ENCODE database (ENCFF022XPK, ENCFF122NPY, ENCFF534EUU, ENCFF846WHK).

### DMR and DMS analysis

Differentially methylated regions (DMRs) were identified using MethScan software[78] with the methscan diff function, employing a bandwidth parameter of 100. Differentially methylated sites were identified using the parameters--stepsize 1 and -- bandwidth 1. The identified DMRs were subsequently merged using BEDTools[70] and clustered via ClusterGvis (https://github.com/junjunlab/ClusterGVis, v0.1.2)

### Motif analysis

For each cell type or clone, de novo motif discovery was performed within differentially methylated regions using the HOMER[79] suite via the findMotifsGenome.pl command, with the parameters-mask and -size 200. Parallel analyses were conducted for nucleosome-depleted regions in each cell type, as well as for differentially accessible regions.

### Epigenetic aging analysis

To investigate methylation age in cancer cells, we utilized the scAge[38] (v1.0) algorithm to estimate the epigenetic age at the single-cell level. The model was trained using 450k methylation array data from LUAD samples sourced from TCGA. Methylation values were binarized to quantify the methylation frequency of WCG sites in each single cell. Statistical concordance (p < 0.05, hypergeometric test) was established between age-associated CG sites and epigenetic clock datasets.

### Enrichment for gene set annotations

Analysis of gene set enrichment was performed using clusterProfiler[80] to elucidate the characteristics of multiomics data. Functional enrichment analysis of peaks was conducted employing cistromego[81]. Bar graph of enriched terms were utilized the web-based platform Metascape (https://metascape.org/gp/index.html#/main/step1).

### Survival analysis

We identified a range of differentially methylated promoter genes and performed comprehensive overall survival analysis for each utilizing the web-based platform GEPIA (http://gepia.cancer-pku.cn).

## Declarations

### Ethics approval and consent to participate

Approval of the study was obtained from the Institutional Review Board of the First Affiliated Hospital of Guangzhou Medical University (Reference number: ES-2025-K007). To investigate the genomic and epigenetic heterogeneity of primary cancer cells using SpliCOOL-seq, we obtained a sample from a patient who had undergone lung resection for pulmonary invasive adenocarcinoma. Written informed consent was provided by the patient before sampling.

### Availability of data and materials

The raw and processed sequencing data in this study have been deposited in the Genome Sequence Archive in National Genomics Data Center, Chinese National Center for Bioinformation/ Beijing Institute of Genomics, Chinese Academy of Sciences (GSA-Human: HRA010465), which are publicly accessible under accession number PRJCA036138. The code for pre-processing SpliCOOL-seq data is accessed online at https://github.com/fanxylab/SpliCOOL-seq.git.

### Consent for publication

The study subject has consented for publication.

### Author’s contributions

X.F. conceived and supervised the project. Q.S. generated the data and E.D. analyzed the data. Q.S. and X.F. wrote the paper. J.Z. cultured the cell lines. Q.Y. prepared the lung cancer sample. D. S., Q.Y., J.Z., E.D. and Q.S. interpreted the data. All authors have read the manuscript, offered feedback and approved it before submission.

## Acknowledgements

We would like to thank the Advanced Cell Technology Core Facility, Guangzhou National Laboratory for offering us the sequencing service in this study.

---

**Figure S1.**
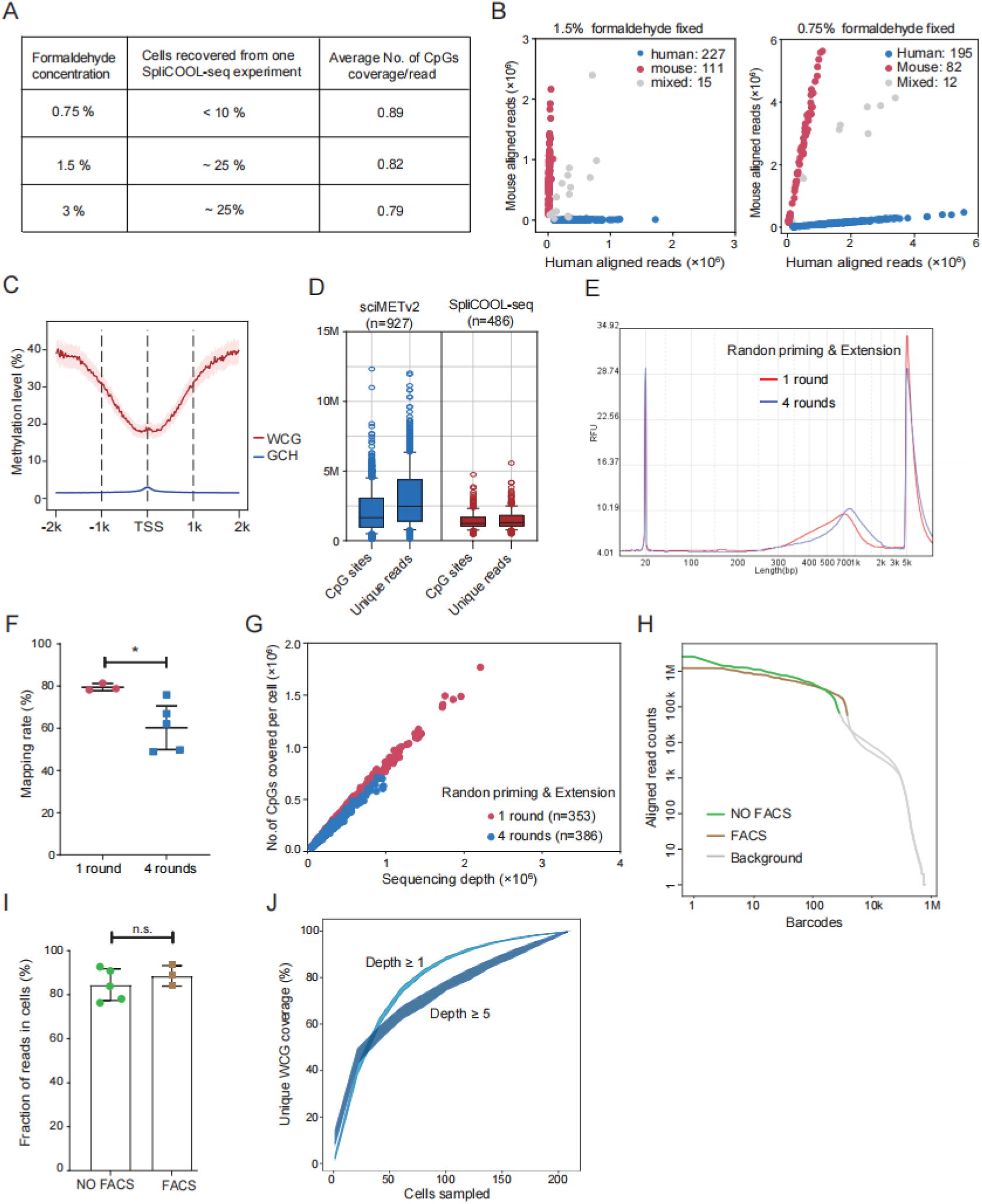
SpliCOOL-seq overview and performance. **A)** Various concentrations of cell fixation have a significant impact on the recovery rate of cells and the sensitivity of CpG site detection. **B)** Scatter plot illustrating the number of reads mapped to the human and mouse genomes in each cell with different concentration of formaldehyde. **C)** WCG and GCH methylation levels of GM12878 cells within ±2 kb of TSS, with shaded regions indicating the 25th and 75th percentiles of methylation levels across cells. M.cviPI treatment with 1.5% formaldehyde fixed cells, cell number n=133. **D)** CpG sites and unique reads in each cell sequenced with sciMETv2 and SpliCOOL-seq. For sciMETv2, CpG sites ranged from 0.5M to 4.5M; unique reads ranged from 0.8M to 6.4M. For SpliCOOL-seq, CpG sites ranged from 0.7M to 2.4M; unique reads ranged from 0.8M to 2.6M. **E)** Fragment size distributions of the libraries with 1 round of random priming and extension and 4 rounds of random priming and extension. **F)** Mapping efficiencies across multiple libraries. (1 round of random priming and extension, n = 3, 4 rounds of random priming and extension, n = 5, mean ±SD). Two-tailed unpaired t test, *p < 0.05. **G** Scatter plot depicting the number of CG dinucleotides covered by the total aligned reads per cell. **H)** Cells ranked based on the count of aligned reads. **I)** Fraction of reads in cells across multiple libraries (experiment without FACS, n=5, with FACS n=3). Two-tailed unpaired t test, n.s., no significance. **J)** Line plots showing the accumulated coverage of WCG sites under different depths with growing number of cells profiled.

**Figure S2.**
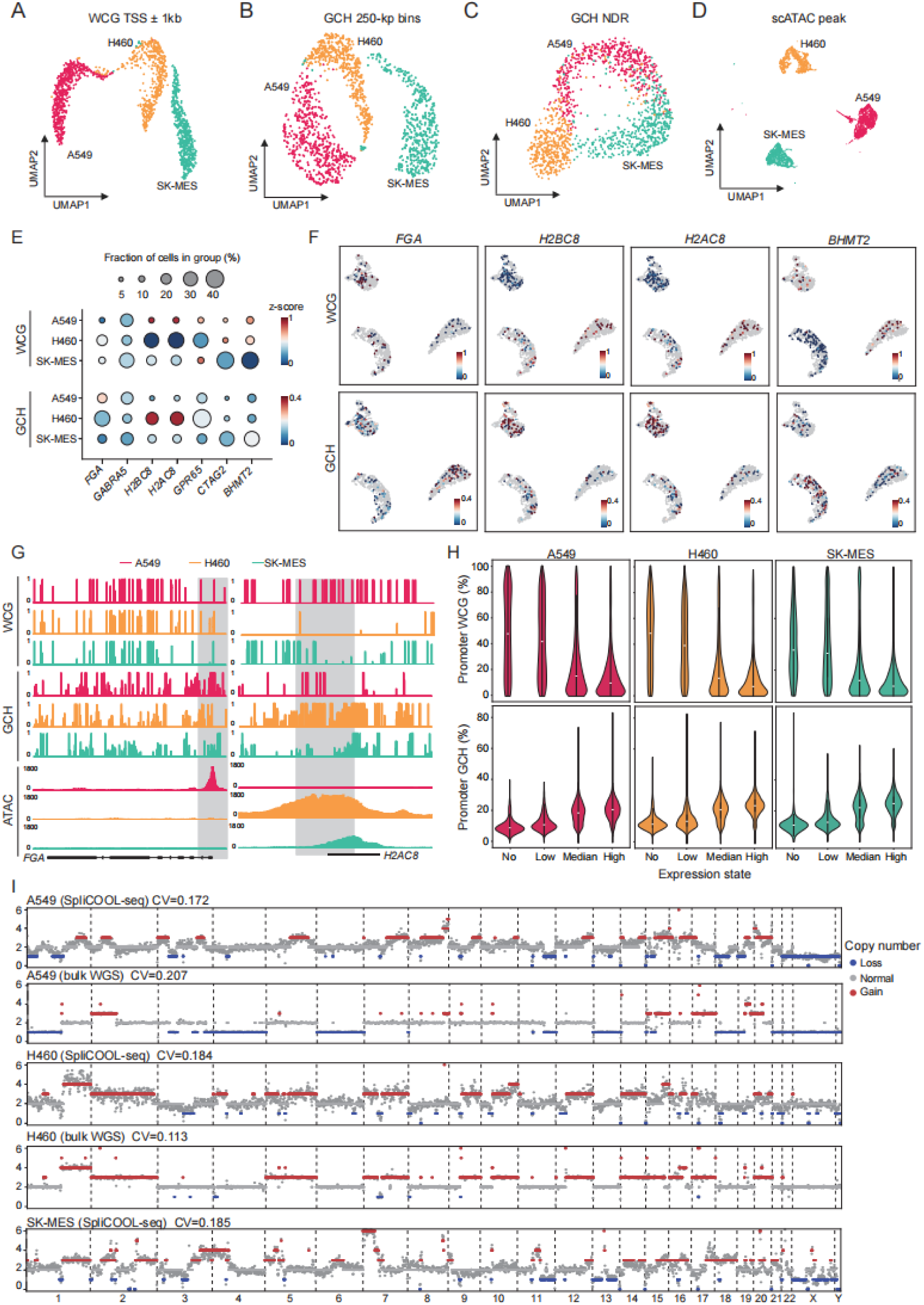
SpliCOOL-seq distinguishes different cell types based on multiple modalities. **A-D)** UMAP embedding showing cells based on their DNA methylation levels (TSS±1kb) (**A**), chromatin accessibility (GCH in 250-kb non-overlapping bins) (**B**), nucleosome depleted region (NDRs)(**C**), and scATAC peak (**D**). Each dot represents an individual cell of A549 cells (red), H460 cells (yellow) and SK-MES cells (blue). **E)** Dot plots showing the promoter WCG methylation levels and GCH methylation levels (−1,000 bp to +1,000 bp) of representative marker genes in the three cell types. **F)** Feature plot showing the DNA methylation levels and chromatin accessibility of each cell type specific genes: A549 (*FGA*), H460 (*H2BC8* and *H2AC8*), and SK-MES (*BHMT2*). Corrected methylation levels are visualized from hypomethylation (blue) to hypermethylation (red). **G)** Browser track showing the DNA methylation (WCG) and chromatin accessibility (GCH or ATAC) profiles at the *FGA* locus and *H2AC8* locus across various cell types. Promoter regions are highlighted by light gray shading. **H)** DNA methylation levels and chromatin accessibility of promoter regions for each gene set categorized by their RNA expression levels within each cell type. No: non-expression; Low: gene expression level is less than 0.25 quantile; Median: Gene expression level is between the 0.25 and 0.75 quantile; High: Gene expression level is above 0.75 quantile. **I)** Normalized signals of read counts and inferred copy numbers of bulk WGS (from ENCODE, see methods) datasets and SpliCOOL-seq (merged single cells) data for each cell type.

**Figure S3.**
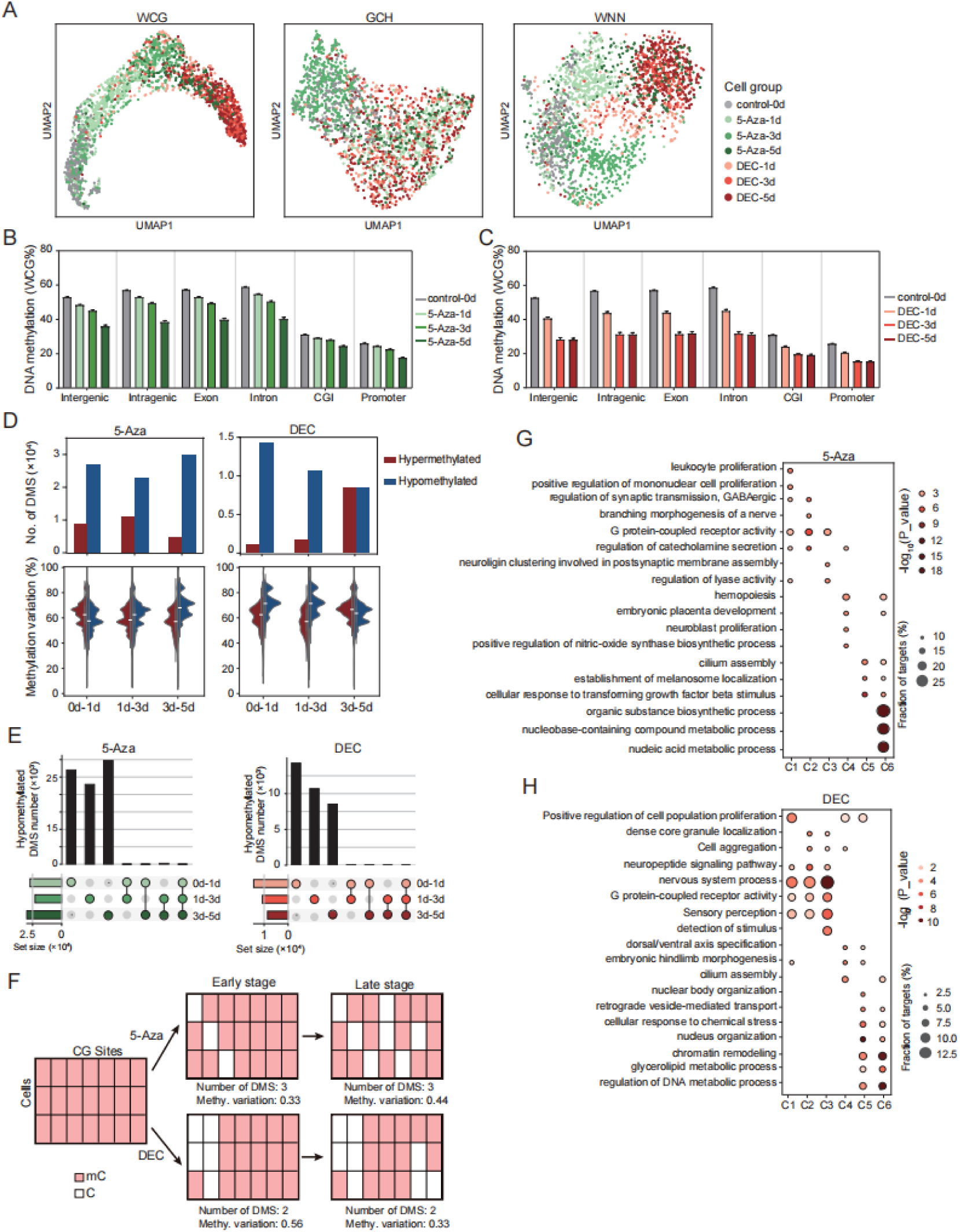
Demethylation patterns of A549 cells treated by different DNMT inhibiters. **A)** UMAP embedding showing cells distributions based on their DNA methylation levels (WCG in 250-kb non-overlapping bins), chromatin accessibility (GCH in 250-kb non-overlapping bins) and an integrated analysis of both modalities. **B, C)** Histogram showing the WCG methylation levels across various genetic elements. **D)** The number (upper panel) and cell proportion (lower panel) of hypermethylated and hypomethylated DMSs between each adjacent timepoints. **E)** Upset plot showing the number of specific and shared hypomethylated DMSs in each stage. **F)** Schematic illustration of the single-cell demethylation process under different treatment. **G, H)** Enriched GO terms for the DMR-associated genes in each cluster.

**Figure S4.**
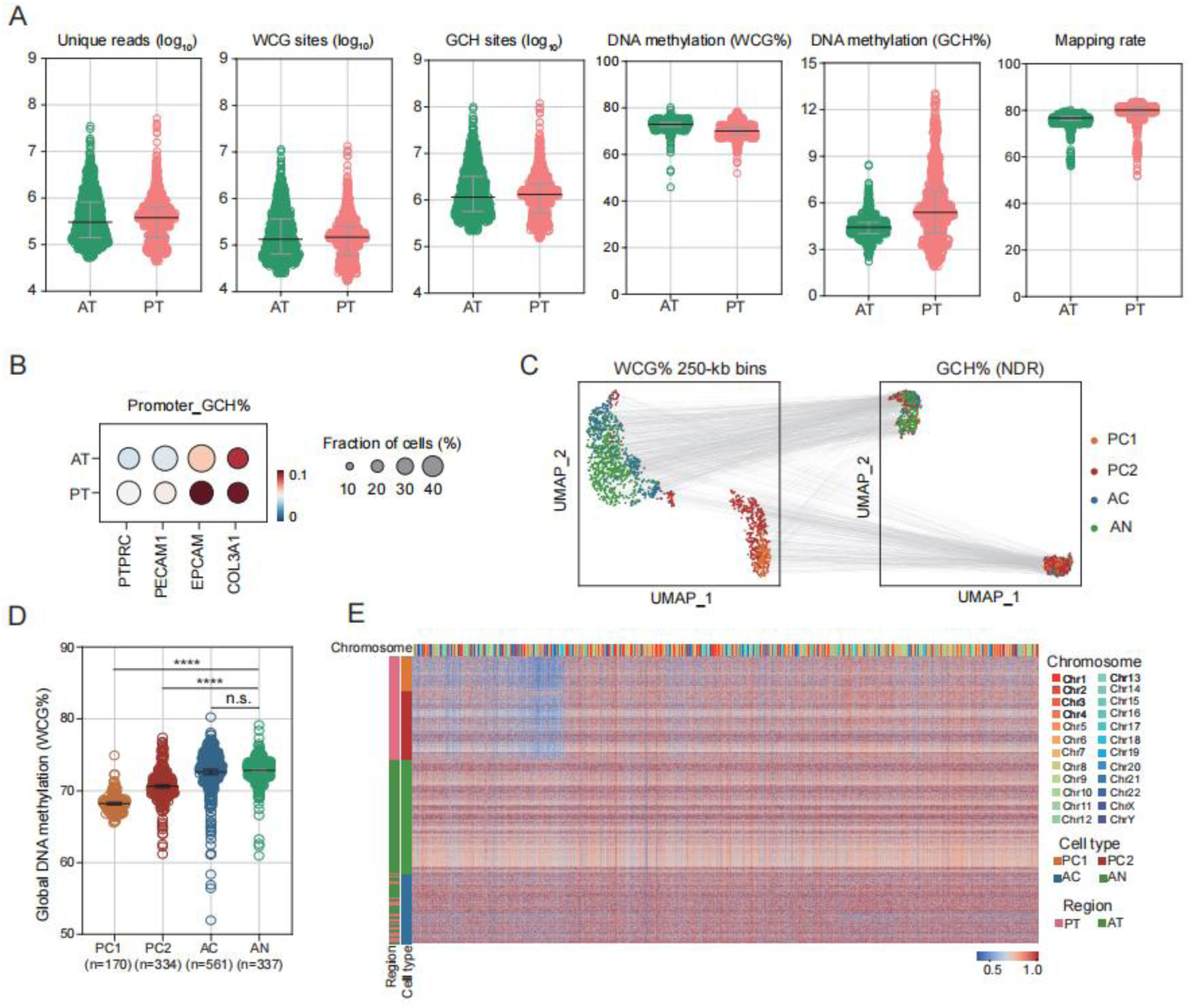
SpliCOOL-seq revealed tumor subclones within the tumor lesion. **A)** Numbers of unique reads, WCG sites and GCH sites covered, global DNA methylation levels, chromatin accessibility, and mapping rate estimated in individual cells from AT and primary PT, respectively. **B)** Dot plots showing the promoter GCH methylation levels (−1,000 bp to +1,000 bp) of representative marker genes in each tissue section. **C)** UMAP embedding illustrating cells based on DNA methylation levels and chromatin accessibility. Each dot represents an individual cell, with connecting lines representing the projection of the same cell. **D)** Global DNA methylation levels of single cells in each clone. Two-tailed unpaired t test, n.s., no significance; ****p < 0.00001. **E)** Heatmap showing the genome-wide DNA methylation levels (10 Mb window) of all cells.

**Figure S5.**
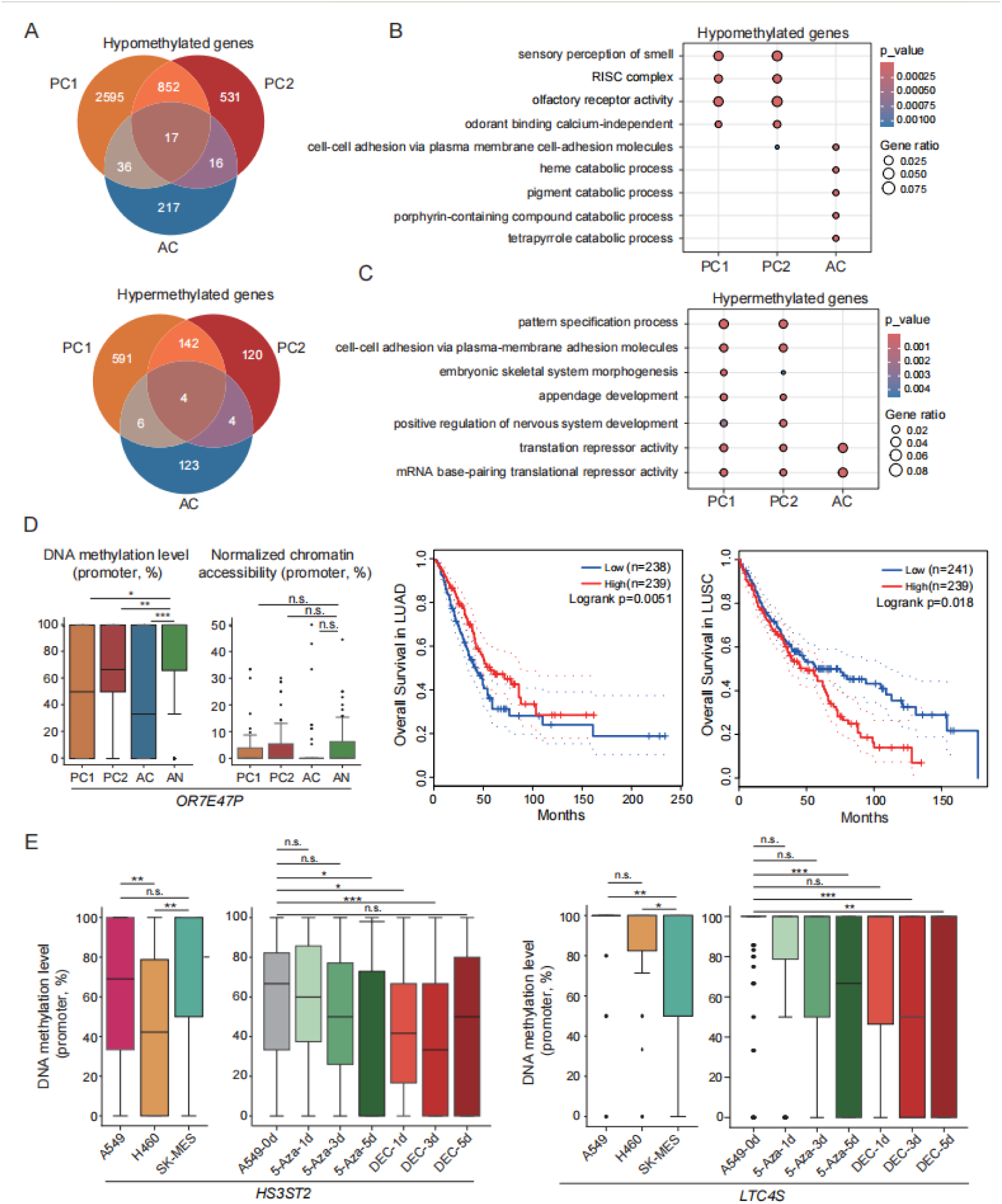
Abnormally methylated genes in the tumor clones. **A)** Venn diagram displaying the intersections of abnormally methylated genes between the three cancerous clones. **B, C)** Enriched GO terms for the DMR-associated genes in each cancerous clone. **D)** DNA methylation levels and chromatin accessibility in the promoter regions of *OR7E47P* across each cell type (left). Overall survival of LUAD and LUSU patients grouped by the expression of *OR7E47P*. **E)** DNA methylation levels in the promoter regions of the two representative genes across different lung cancer cell lines (left) and in DNMT inhibitor treated A549 cells (right). The statistical test was carried out using the Wilcoxon rank-sum test. n.s., no significance; *P < 0.05; **P < 0.01; ***P < 0.001.

**Figure S6.**
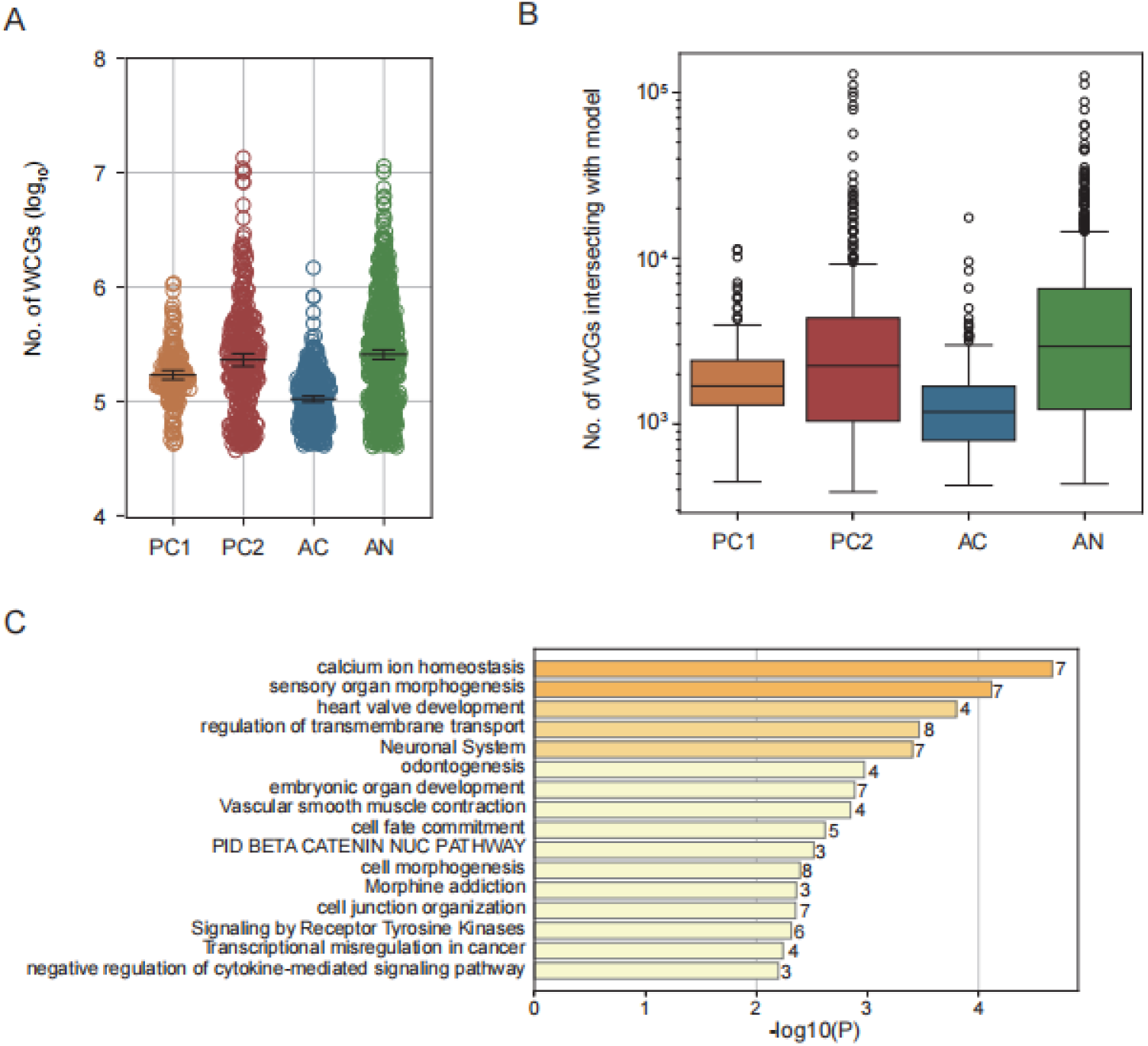
Consistency of aging and tumor related DNA methylation changes. **A)** Numbers of WCG sites covered in individual cells of each clone. **B)** Boxplot showing numbers of WCG sites intersecting with the 450K chip. **C)** Bar graph of enriched terms for the predictive sites which also showed abnormal methylation in cancer cells, colored by p-values, number of genes on the right.

